# Distinct immune profiles in children living with HIV based on timing and duration of suppressive antiretroviral treatment

**DOI:** 10.1101/2024.08.27.609833

**Authors:** Madeline J Lee, Morgan L Litchford, Elena Vendrame, Rosemary Vergara, Thanmayi Ranganath, Carolyn S Fish, Daisy Chebet, Agnes Langat, Caren Mburu, Jillian Neary, Sarah Benki, Dalton Wamalwa, Grace John-Stewart, Dara A Lehman, Catherine A Blish

## Abstract

Timely initiation of antiretroviral therapy (ART) remains a major challenge in the effort to treat children living with HIV (“CLH”) and little is known regarding the dynamics of immune normalization following ART in CLH with varying times to and durations of ART. Here, we leveraged two cohorts of virally-suppressed CLH from Nairobi, Kenya to examine differences in the peripheral immune systems between two cohorts of age-matched children (to control for immune changes with age): one group which initiated ART during early HIV infection and had been on ART for 5-6 years at evaluation (early, long-term treated; “ELT” cohort), and one group which initiated ART later and had been on ART for approximately 9 months at evaluation (delayed, short-term treated; “DST” cohort). We profiled PBMC and purified NK cells from these two cohorts by mass cytometry time-of-flight (CyTOF). Although both groups of CLH had undetectable viral RNA load at evaluation, there were marked differences in both immune composition and immune phenotype between the ELT cohort and the DST cohort. DST donors had reduced CD4 T cell percentages, decreased naive to effector memory T cell ratios, and markedly higher expression of stress-induced markers. Conversely, ELT donors had higher naive to effector memory T cell ratios, low expression of stress-induced markers, and increased expression of markers associated with an effective antiviral response and resolution of inflammation. Collectively, our results demonstrate key differences in the immune systems of virally-suppressed CLH with different ages at ART initiation and durations of treatment and provide further rationale for emphasizing early onset of ART.

**AUTHOR SUMMARY:** Many children living with HIV lack access to both antiviral treatments and testing for HIV infection and are therefore unable to initiate treatment in a timely manner. When children do begin treatment, their immune systems take time to recover from the uncontrolled HIV infection. In this study, we examine how the immune systems of children living with HIV normalize after treatment onset by looking at two groups of children whose HIV is well-controlled by treatment and who therefore don’t have virus replicating in their blood. One group started treatment within the first year of life and has been on treatment for 5-6 years, while the other began treatment after the first year and has been treated for around 9 months. Although both of these groups are virally-suppressed, we found significant differences in their immune profiles, with the children who had delayed and short-term treatment showing signs of inflammation and immune dysfunction. Collectively, our study helps us understand how variation in the timing and duration of ART treatment impacts the immune system in children with viral suppression and therefore provides clinicians with additional knowledge that can inform the care of children living with HIV, improving their health and quality of life.

## INTRODUCTION

It is estimated that around 1.4 million children worldwide are currently living with HIV. The increasing availability of antiretroviral therapy (ART) has contributed to a significant decrease in mortality among children living with HIV (CLH) and the percentage of CLH receiving ART has increased from roughly 18% worldwide in 2010 to nearly 60% in 2022 (https://data.unicef.org/topic/hivaids/paediatric-treatment-and-care/). However, several major problems remain in the treatment of CLH, including continuing access to medication and timely initiation of ART. The majority of HIV infections acquired by children occur via perinatal transmission(1,2). Children who initiate ART earlier have smaller HIV reservoir sizes, longer time to viral rebound following ART interruption, decreased immune activation at baseline, and decreased overall mortality(3–9). Despite the known benefits of early ART intervention, many CLH are undiagnosed and hence unable to initiate ART during acute infection.

ART attenuates key aspects of immune dysregulation that result from HIV-1 infection, including CD4 T cell loss and CD8 T cell activation in both adults and children(10–12). However, some of these factors never completely normalize, particularly in individuals with later initiation of ART. In 2017, Alvarez et al. conducted one of the only studies to examine the kinetics of immune normalization in ART-treated CLH; they found that CD4 loss and CD8 T cell activation in CLH are reversed to levels seen in children without HIV within 10 months of ART initiation(10). While this study provided valuable insight, it examined only a few metrics of immune restoration following ART initiation in CLH.

In this study, we analyze the temporal dynamics of immune normalization following ART initiation in CLH. We leverage peripheral blood samples from CLH with viral suppression from two cohorts from early in the ART era: one which initiated ART during acute/early HIV infection and one which initiated ART later, during chronic infection. In order to minimize developmental differences in the immune systems of our two cohorts, we utilized samples which were collected at similar ages across both cohorts (median = 6.08 years), at which point the early-treated children had been on ART for 5-6 years (early long-term treated, ELT) while the delayed-treated children had been on ART for approximately 9 months (delayed, short-term treated, DST). We use these samples to demonstrate that, although the children with delayed, short-term treatment had undetectable viral loads, they had severely altered peripheral immune systems compared to the children who had early, long-term treatment. The DST children had immune systems marked by altered T cell memory subset frequencies and upregulation of stress-induced surface proteins. Conversely, the ELT children exhibited increased expression of markers associated with lymphocyte maturity and a productive antiviral response. Collectively, our study demonstrates that early treatment and longer duration of ART has a marked impact on immune recovery. Thus, two age-matched children (eg., both 5 years old) with perinatal HIV infection and current viral suppression may have markedly different immune phenotypes due to differences in timing and duration of ART.

### Cohort

In this study, we sought to understand how the duration of untreated HIV and timing of treatment initiation influence the pediatric immune landscape, which changes rapidly in the first few years of life(13–15). To do this, we profiled the peripheral immune cells from two cohorts of ART-treated CLH in Nairobi, Kenya between 2005-2007, a time prior to the current recommendation of early ART initiation (**Fig. 1A**): one cohort in which children initiated ART soon after birth (the early long-term treatment “ELT” cohort); and one in which children were diagnosed with HIV and initiated ART later in life (the delayed short term-treatment “DST” cohort) (**Table 1**). To account for the age-dependent changes in immune profiles, PBMC samples collected from children at similar age ranges (between 4 and 8 years of age) were included from both cohorts, corresponding to approximately 9 months post-ART initiation for DST donors (n=19) and 66 months after ART initiation for ELT donors (n=25) (**Fig. 1C**). As our goal was to assess immune status in children who had fully suppressed their viral load, we only included children who had a viral load below 150 copies/mL (the assay’s limit of detection) for >3 months prior to immune status assessment as well as at the time of sample collection. Consistent with the later initiation of ART, the DST cohort samples had a slightly higher level of total T cell-associated HIV DNA, indicating a larger viral reservoir, although their levels of intact T cell-associated HIV DNA were not significantly different (**Fig. 1D**).

**Figure 1:**
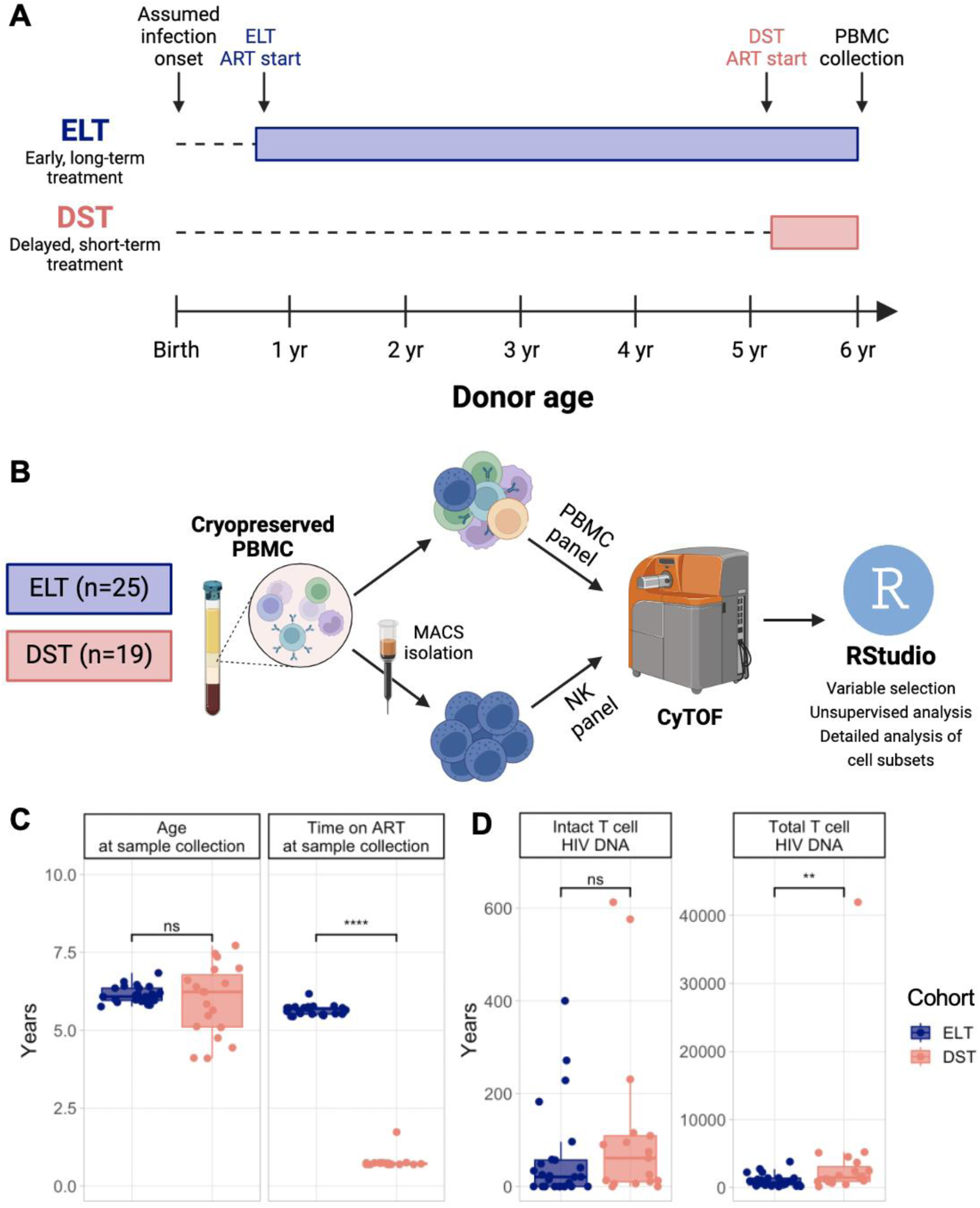
Overview of cohorts. A) Schematic showing the age and duration of ART for the two cohorts used in this study. Blue and pink bars indicate the duration of ART for the early, long-term treatment cohort (ELT; top, blue) and the delayed, short term treatment cohort (DST; bottom, pink). Dashed lines indicate the duration of untreated HIV-1 infection. All donors were assumed to be infected before, at, or near birth. ELT donors initiated ART within the first year of life. DST donors initiated ART at ages 3-7. B) Schematic illustration of experimental design. C) Boxplots showing the age of each donor at PBMC collection (left) and the length of time each donor had been on ART at sample collection (right). D) Boxplots showing the number of copies of intact T cell-associated HIV DNA (left) and total T cell-associated HIV DNA (right) for each donor. HIV DNA levels were determined by cross subtype-intact proviral DNA assay (CS-IPDA). Significance values were determined using a Wilcoxon rank-sum test. ns, p > 0.05. *, p < 0.05. **, p < 0.005. ***, p < 0.0005. ****, p < 0.00005.

**Table 1:** Key demographic and clinical information for donors used in this study. “Undetectable” refers to viral loads below the assay’s limit of detection, which in this case was 150 copies/mL of viral RNA. All donors in this study had undetectable viral loads.

|  | ELT | DST | Overall |
| --- | --- | --- | --- |
|  | (N=25) | (N=19) | (N=44) |
| **Sex** |  |  |  |
| Male | 7 (28.0%) | 9 (47.4%) | 16 (36.4%) |
| Female | 18 (72.0%) | 10 (52.6%) | 28 (63.6%) |
| **Age (years)** |  |  |  |
| Mean (SD) | 6.13 (0.265) | 5.95 (1.13) | 6.05 (0.762) |
| Median [Min, Max] | 6.08 [5.76, 6.84] | 6.23 [4.10, 7.72] | 6.09 [4.10, 7.72] |
| **Age at ART onset (years)** |  |  |  |
| Mean (SD) | 0.486 (0.186) | 5.18 (1.14) | 2.51 (2.47) |
| Median [Min, Max] | 0.410 [0.210, 0.830] | 5.15 [3.36, 6.98] | 0.755 [0.210, 6.98] |
| **Time on ART (years)** |  |  |  |
| Mean (SD) | 5.65 (0.155) | 0.767 (0.234) | 3.54 (2.45) |
| Median [Min, Max] | 5.69 [5.44, 6.17] | 0.710 [0.690, 1.73] | 5.49 [0.690, 6.17] |
| **Viral load** |  |  |  |
| **(RNA copies/mL)** |  |  |  |
| Mean (SD) | Undetectable | Undetectable | Undetectable |
| Median [Min, Max] | Undetectable | Undetectable | Undetectable |

## RESULTS

### The immune profiles of early, long-term treated children are distinct from those of delayed, short term-treated children

We performed mass cytometry by time-of-flight (CyTOF) on whole peripheral blood mononuclear cells (PBMC) and purified natural killer (NK) cells from the two pediatric cohorts and first sought to characterize broad changes in the immune composition and phenotype between the two groups. We stained each sample with two panels of antibodies conjugated to heavy metals (one for whole PBMC and one for purified NK cells); these panels, which have been previously described in detail, are designed to deeply interrogate the immune response to viral infection with a focus on ligand/receptor interactions between NK cells and other peripheral immune cells (16). Although both ELT and DST cohorts were virally suppressed at the time of sample collection, there were marked differences in the peripheral immune profiles between the two cohorts. After nine months of ART treatment and at least three months of viral suppression, DST children had a significantly lower frequency of CD4 T cells and correspondingly higher CD8 T cell levels than ELT children that had been on ART for ∼5-6 years (**Fig. 2A**). The overall frequency of T cells, as well as B cells, monocytes, and NK cells, was unchanged between the two cohorts (**Fig. 2A**).

**Figure 2:**
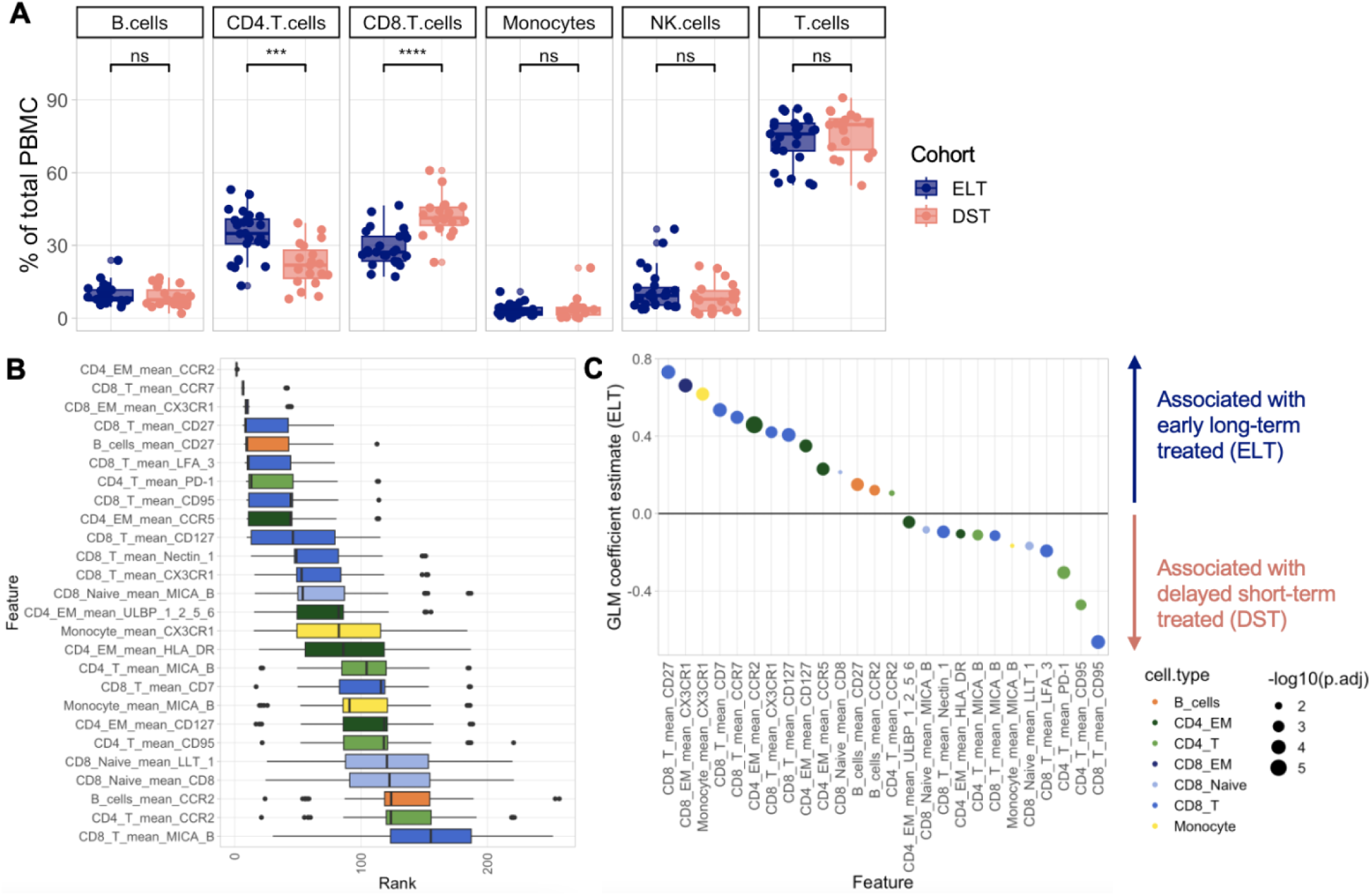
Age-matched CLH in the ELT and DST cohorts exhibit broad differences in immune composition and phenotype. A) Boxplots showing the frequencies of major immune cell subsets as a percentage of total PBMC in each cohort. B) Boxplot showing the variables selected by MUVR. The X axis shows the rank of the importance of each variable in the model in each iteration of the analysis, with a lower rank being more important. Boxes are colored by cell type. C) GLM coefficient estimates of each variable selected in (B) with the ELT cohort. Positive coefficient values indicate an association with the ELT cohort; negative coefficient values indicate an association with the DST cohort. Dot color indicates cell type. Dot size indicates -log10 adjusted P value; P values were adjusted with the Benjamini-Hochberg correction for multiple hypothesis testing. Variables selected in (B) with an adjusted P value > 0.05 were not plotted in C. Significance values were determined using a Wilcoxon rank-sum test. ns, p > 0.05. *, p < 0.05. **, p < 0.005. ***, p < 0.0005. ****, p < 0.00005.

We next interrogated changes in immune phenotype between the cohorts using **<u>MU</u>**ltivariate modeling with minimally biased **<u>V</u>**ariable selection in **<u>R</u>** (MUVR), a random forest-based method of determining which variables are the most important in determining an outcome variable(17). A total of 287 continuous variables were used for MUVR analysis (**Supplemental Table 1**). These included mean expression of each marker in our CyTOF panels in each of 9 cell types (B cells, monocytes, total CD4 T cells, naive CD4 T cells, effector memory CD4 T cells, total CD8 T cells, naive CD8 T cells, effector memory CD8 T cells, and NK cells). The outcome variable was binary classification by cohort (ELT or DST). This analysis identified 26 predictor variables, representing all cell types except naive CD4 T cells and NK cells (**Fig. 2B**). We then utilized a generalized linear model (GLM) to stratify these predictor variables by their association with either the ELT or DST cohorts (**Fig. 2C**). 14 variables were significantly associated with the ELT cohort, while 12 variables were significantly associated with the DST cohort. The 12 variables whose expression were associated with the DST cohort were markers of activation and exhaustion (**Fig. 2C**). These markers are MICA/B, Nectin-1, HLA-DR, LFA-3, PD-1, ULBP, and CD95 in T cells and MICA/B expression in monocytes. Conversely, the T cell markers (CD27, CX3CR1, CD7, CCR7, CCR2, CD127, and CCR5) associated with ELT cohort donors tended to be associated with survival, terminal differentiation, and antiviral response (**Fig. 2C**). ELT cohort donors also had higher CX3CR1 expression in monocytes (**Fig. 2C**). Collectively, these results suggest that there exist differences in immune composition and phenotype between early long-term- and delayed short-term-treated donors, with the ELT donors exhibiting upregulation of markers involved with a healthy and productive antiviral immune response and DST donors expressing higher levels of stress-associated markers. These differences may impact these donors’ ability to respond to infection, malignancy, or other challenges to the immune system, as DST donors appear to have a much higher level of background inflammation and a dearth of CD4 T cells.

### NK cells appear less mature and less functional in DST children compared to those of ELT children

Having identified overall changes in the immune composition and phenotype of children in the DST cohort compared to those in the ELT cohort, we interrogated changes within individual immune cell subsets, beginning with NK cells as our NK cell-specific CyTOF panel allowed us to identify granular changes in NK cell composition and phenotype. We began by analyzing the frequencies of four major NK cell subsets: CD56^bright^ NK cells, CD56^dim^CD16^hi^ NK cells, CD56^dim^CD16^lo^ NK cells, and CD56-NK cells. We found that DST donors had a significantly lower percentage of CD56^dim^CD16^hi^ NK cells, which are typically mature, cytotoxic NK cells. These donors also had trending increases in the abundances of CD56^bright^CD16^low^ NK cells, which are immature and produce high levels of cytokines; CD56^dim^CD16^low^ NK cells, an unconventional mature NK cell population with reduced antibody-dependent cellular cytotoxicity (ADCC) capacity; and CD56^-^CD16^hi^ NK cells, which are a dysfunctional population that expand during chronic infection (**Fig. 3A**).

**Figure 3:**
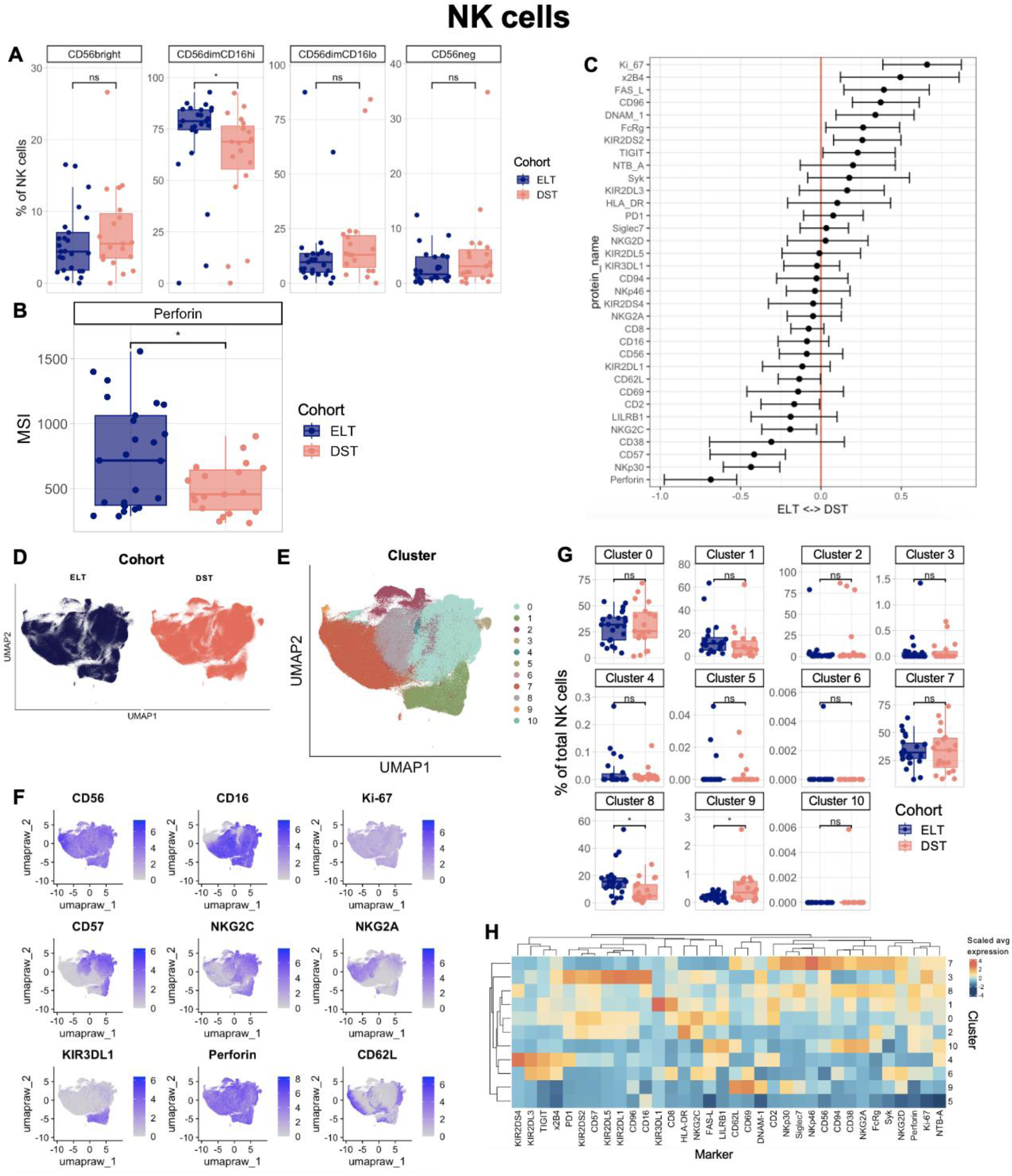
DST donor NK cells appear immature and less functional compared to those of ELT children. A) Boxplots showing the frequencies of major NK cell subsets as a percentage of all NK cells in the two cohorts. CD56^bright^ NK cells are defined as CD56^bright^, CD16^lo^. B) Boxplots showing the median signal intensity (MSI) in all NK cells of three markers that were differentially expressed at the patient level between ELT and DST donors. C) CytoGLMM analysis of NK features that are significant predictors of the ELT cohort (left) and the DST cohort (right). Each row represents one marker. The X axis represents log-odds. Bars represent 95% confidence intervals. Markers whose 95% confidence intervals do not cross the red line are considered to be significant predictors of one cohort over the other. D-E) UMAP embeddings of the NK cells in the dataset, colored by cohort (D) or PARC cluster (E). F) Feature plots showing the expression of 9 different proteins on the NK cells in this dataset. G) Boxplots showing the frequency of each PARC cluster among all NK cells in the dataset. H) Heatmap showing the scaled expression of each marker in each PARC cluster. Significance values were determined using a Wilcoxon rank-sum test. ns, p > 0.05. *, p < 0.05. **, p < 0.005. ***, p < 0.0005. ****, p < 0.00005.

Two markers were differentially expressed on bulk NK cells between the two cohorts: CD57, a marker of maturity; and CD62L, which denotes polyfunctional and intermediately mature NK cells(18)(**Fig. 3B**). Both of these markers were upregulated in ELT donors compared to DST donors. To identify additional differences between the cohorts, we used a generalized linear mixed model to find markers whose expression was associated with one cohort over the other(19). In addition to CD57, and CD62L, ELT donor status was correlated with expression of the cytotoxic molecule Perforin as well as the activating receptors NKp30, CD16, and CD2 ( **Fig. 3C**). Meanwhile, DST donor status was correlated with expression of all three members of the DNAM-1/TIGIT/CD96 axis, which regulates NK cell activation and inhibition via ligation of Nectin-2 and poliovirus receptor(20). Higher levels of proliferation marker Ki-67, the apoptosis inducer Fas-L, and the marker of decreased NK cell functionality FcRg were also associated with the DST cohort (**Fig. 3C**).

To further interrogate changes in NK cell phenotype, we generated a UMAP embedding of the NK cells in our dataset (**Fig. 3D-F**). As suggested by our previous findings, we observed a high degree of overlap in the distribution of the NK cells from the two individual cohorts (**Fig. 3D**). However, when we performed unsupervised clustering on the NK cells using the PARC algorithm(21), we did identify two clusters that were differentially abundant between DST and ELT donors (**Fig. 3E,G**). Cluster 8 is a large cluster (typically comprising 5-20% of peripheral NK cells) that was more abundant in ELT donors (**Fig. 3E,G**) and exhibited an intermediately-mature phenotype that falls between the more mature (CD57^hi^ CD56^dim^ CD16^hi^) NK cells and the less mature (CD57^low^ CD56^bright^ CD16^low^) NK cells (**Fig. 3E-F**). This cluster also had the highest Perforin expression out of any of the clusters (**Fig. 3H)**. Cluster 9 was the only cluster more abundant in DST donors. This is a much smaller cluster, comprising only 0-3% of all NK cells (**Fig. 3G**), and is an unconventional CD56^dim^ CD16^low^ population with very high expression of CD62L and CD69 and particularly low expression of Perforin (**Fig. 3H**). Overall, these analyses suggest that ELT donor NK cells were broadly more mature and functional compared to DST donor NK cells, which were more proliferative but less cytotoxically capable.

### DST children have an increased frequency of nonclassical and stressed monocytes compared to ELT children

We next analyzed changes in the composition and phenotype of monocytes across CLH with delayed, short-term-treatment (DST) and early, long-term treatment (ELT). Monocytes from donors in the ELT cohort expressed higher levels of the chemokine receptor CX3CR1, which is associated with homeostasis(22,23) (**Fig. 4A**). Meanwhile, DST donor monocytes expressed comparatively more LLT-1, MICA/B, and PD-1, all of which can be upregulated in the context of infection and cellular stress(24,25) (**Fig. 4A**). Further analysis by generalized linear mixed model (GLMM) found that expression of the adhesion molecule LFA-3, HLA-C, and HLA-Bw4 was significantly associated with the ELT cohort, while CCR7 and HLA-Bw6 were associated with the DST cohort (**Fig. 4B**). Collectively, these findings suggest that monocytes from patients with delayed and shorter-term treatment are under significant stress and downregulate markers associated with homeostasis and inflammation resolution in comparison to early, long-term-treated donors.

**Figure 4.**
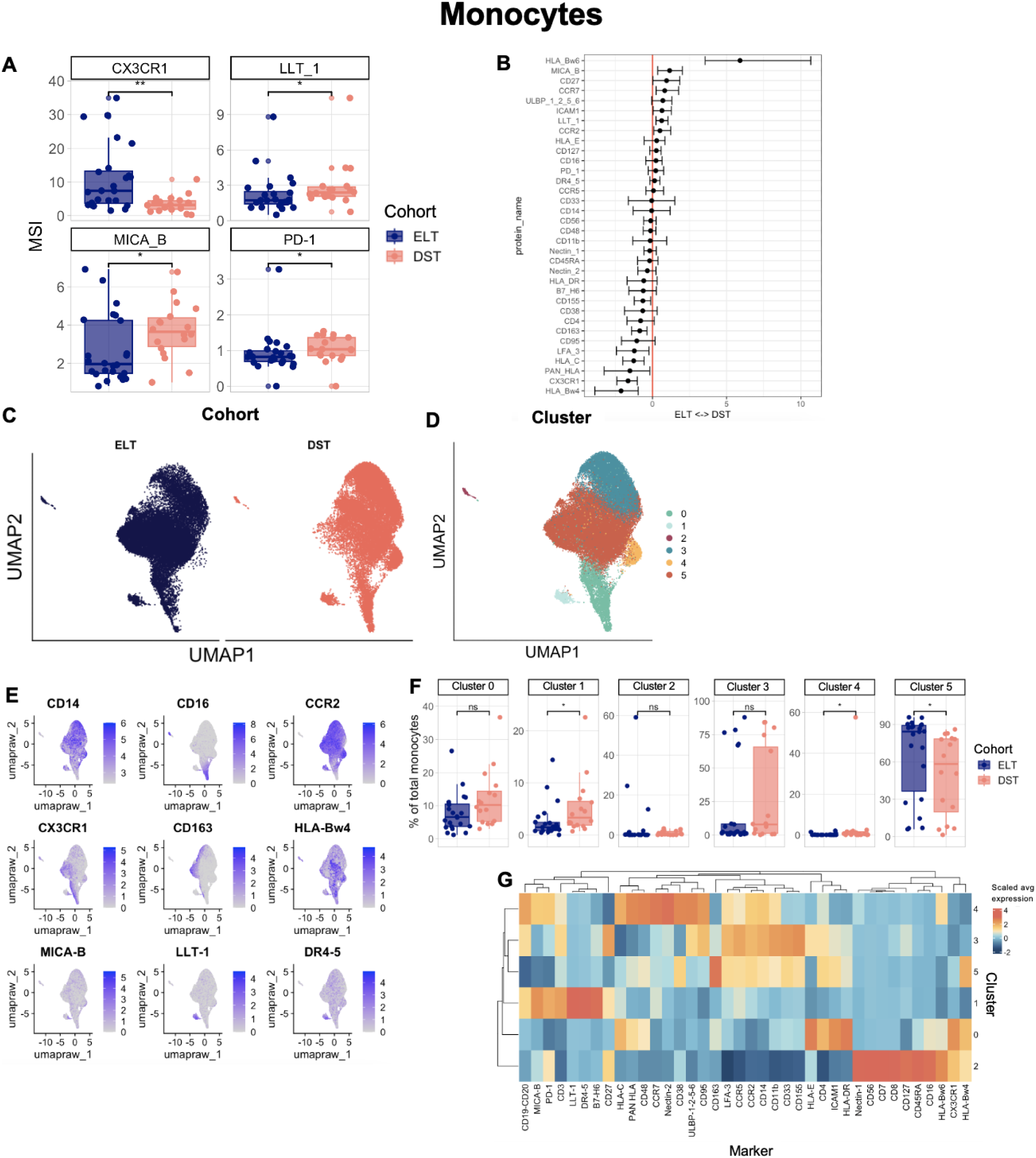
A) Boxplots showing the median signal intensity (MSI) in all monocytes of four markers that were differentially expressed at the patient level between ELT and DST donors. B) CytoGLMM analysis of monocyte features that are significant predictors of the ELT cohort (left) and the DST cohort (right). Each row represents one marker. The X axis represents log-odds. Bars represent 95% confidence intervals. Markers whose 95% confidence intervals do not cross the red line are considered to be significant predictors of one cohort over the other. C) UMAP embeddings of the monocytes in the dataset, colored by cohort (C) or PARC cluster (D). E) Feature plots showing the expression of 6 different proteins on the monocytes in this dataset. F) Boxplots showing the frequency of each PARC cluster among all monocytes in the dataset. G) Heatmap showing the scaled expression of each marker in each PARC cluster. Significance values were determined using a Wilcoxon rank-sum test. ns, p > 0.05. *, p < 0.05. **, p < 0.005. ***, p < 0.0005. ****, p < 0.00005.

Unsupervised analysis of the monocytes in the ELT and DST cohorts also revealed differences between the delayed short-term- and early long-term-treated donors. A UMAP embedding of all monocytes in the dataset illustrates more significant shifts between the two cohorts than was observed in NK cells (**Fig. 4C**). Notably, DST donor monocytes were overrepresented in the portions of the UMAP embedding with high CD16 expression, indicating a higher abundance of nonclassical monocytes in this cohort (**Fig. 4C-E**). When we performed unsupervised clustering on our monocyte dataset, we did indeed find a cluster with high CD16 expression (cluster 0) that was more abundant in DST donors than ELT donors, although this difference is not statistically significant (**Fig. 4F-G**). Another cluster, cluster 1, was present at a significantly higher frequency in DST donors; this cluster was defined by high expression of several stress-induced proteins (MICA/B, LLT-1, B7-H6, and DR4/5) as well as PD-1 (**Fig. 4F-G**). Finally, cluster 5, which had particularly high expression of CD163 and HLA-Bw4, was more abundant in ELT donors (**Fig. 4F-G**). Overall, these results illustrate the shift towards nonclassical and stressed monocytes in CLH with delayed, short-term treatment.

### CD4 T cell memory subsets are significantly altered in ELT children compared to DST children

In earlier analyses, we found that CD4 T cell frequency was lower in DST children, despite these children having HIV RNA measured below detection for at least 3 months prior to sample collection (**Fig. 2A**). Delving into a more granular CD4 T cell analysis, we identified striking differences in the memory T cell subset composition of both CD4 T cells between the cohorts: DST donors had a significantly lower abundance of naive CD4 T cells and a proportionally higher abundance of effector memory (EM) and effector memory re-expressing CD45RA (EMRA) T cells, which are a terminally-differentiated subset of effector memory T cells(26) (**Fig. 5A**). The overall phenotype of CD4 T cells was also significantly different in DST donors compared to the ELT cohort, with stress-induced markers like LLT-1, PD-1, and CD95 being associated with DST donors (**Fig. 5B**). CCR2, CD48 and CX3CR1 were all significantly associated with ELT (**Fig. 5B**).

**Figure 5.**
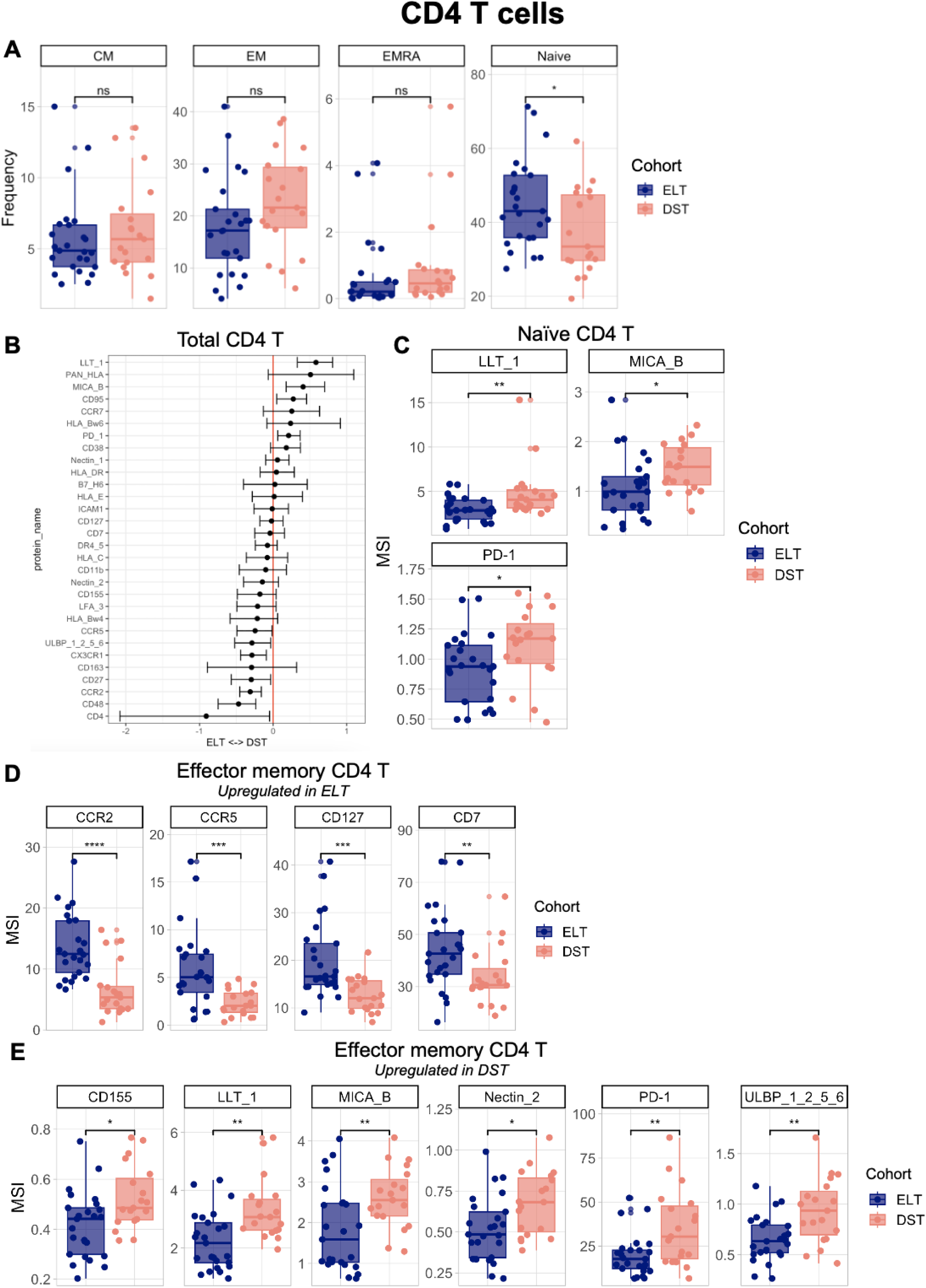
A) Boxplots showing the frequencies of memory CD4 T cell subsets as a percentage of total CD4 T cells in each cohort. B) CytoGLMM analysis of total CD4 T cell features that are significant predictors of the ELT cohort (left) and the DST cohort (right). Each row represents one marker. The X axis represents log-odds. Bars represent 95% confidence intervals. Markers whose 95% confidence intervals do not cross the red line are considered to be significant predictors of one cohort over the other. C-E) Boxplots showing the median signal intensity (MSI) in naive CD4 T cells (C) or effector memory CD4 T cells (D-E) four markers that were differentially expressed at the patient level between ELT and DST donors. Markers shown in D are expressed at higher levels in ELT patients; markers shown in E are expressed at higher levels in DST patients. Significance values were determined using a Wilcoxon rank-sum test. ns, p > 0.05. *, p < 0.05. **, p < 0.005. ***, p < 0.0005. ****, p < 0.00005.

Given that some of these shifts in bulk T cell phenotype may reflect the differing T cell memory subset composition of the two cohorts, we next interrogated phenotypic shifts in the two largest CD4 T cell subsets, naive and effector memory CD4 T cells. Naive CD4 T cells were the subset with the fewest changes between the two cohorts, with DST donor naive CD4 T cells only expressing higher levels of LLT-1, MICA/B, and PD-1 (**Fig. 5C**). Effector memory (EM) CD4 T cells, however, were substantially different between the two cohorts. CD4 TEM cells in ELT donors had significantly higher expression of both CCR2 and CCR5, which collectively mark type 1 helper T cells (Th1 cells)(27). CCR5 can also act as a co-receptor for HIV entry(28); the lower levels of CCR5 and CD4 expression in the CD4 TEM cells of DST donors may reflect the fact that more of these cells are recently infected/depleted and not yet reconstituted. CD4 TEM cells of ELT patients also have increased CD127 expression, which is a marker of successful immune recovery in HIV-1 infection(29) (**Fig. 5D**). Additionally, as observed in other cell subsets, CD4 TEM cells from DST cohort samples had significantly higher expression of a variety of stress-induced molecules (**Fig. 5E**).

### CD8 T cell memory subsets are significantly altered in ELT children compared to DST children

Finally, we analyzed CD8 T cell subset distribution and phenotype in our delayed short-term-treated (DST) cohort compared to our early long-term-treated (ELT) cohort. Similar to CD4 T cells, we found that CD8 T cell distribution was skewed towards effector memory subsets in DST donors, whereas ELT donors had a higher relative abundance of naive CD8 T cells (**Fig. 6A**). The proportion of central memory CD8 T cells and NKT cells was unchanged (**Fig. 6A**). The overall CD8 T cell pool phenotype was likewise disrupted in the DST cohort; like CD4 T cells, the CD8 T cells of these donors had higher expression of stress-induced proteins like LLT-1, MICA/B, CD95, CD38, and ULBPs 1, 2, 5, and 6. ELT CD8 T cells were associated with higher expression of CX3CR1, CD8, CD27, CCR2, and CD48. Interestingly, increased HLA-DR expression, which typically marks CD8 T cell activation, was also associated with the ELT cohort (**Fig. 6B**).

**Figure 6.**
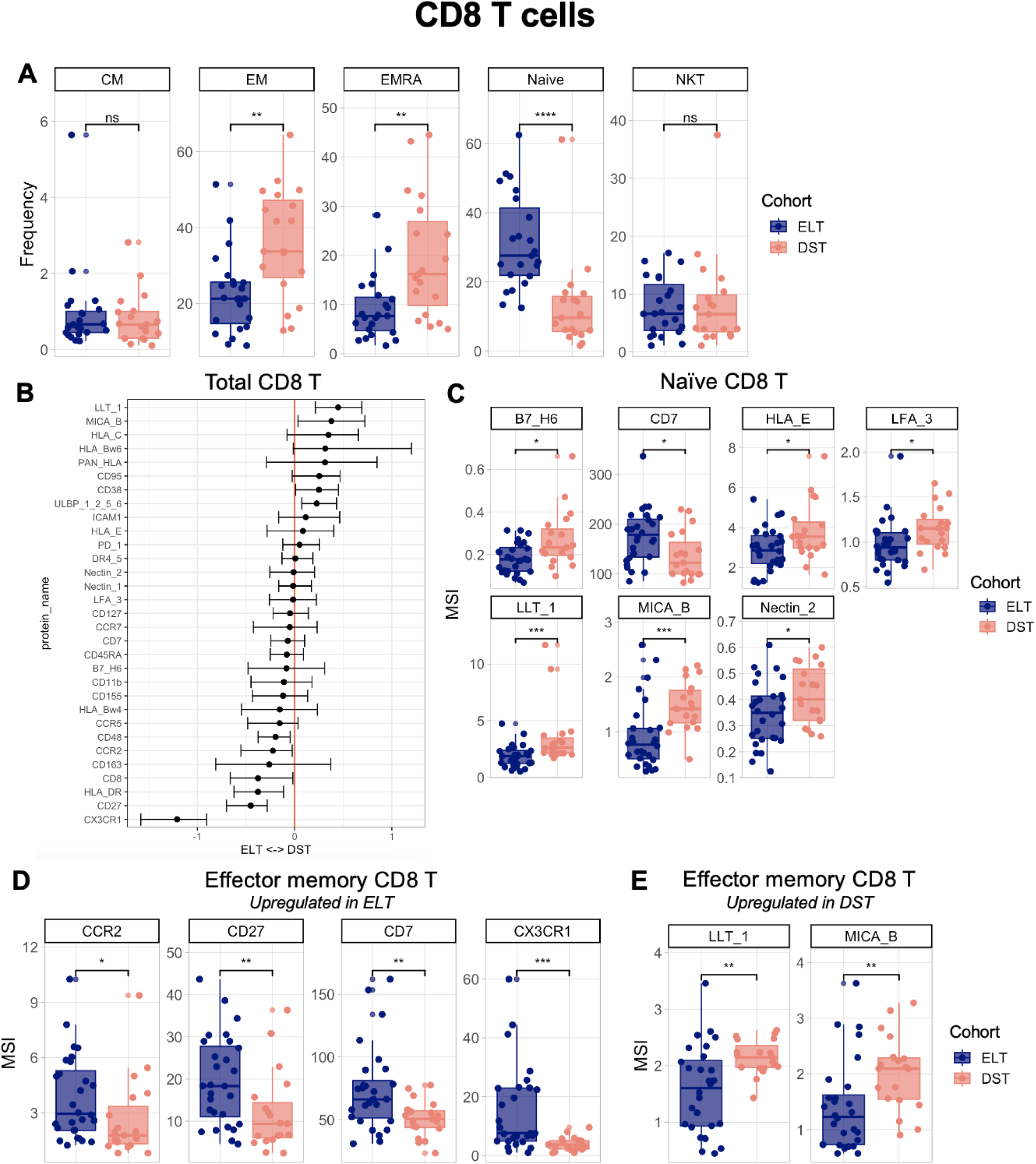
A) Boxplots showing the frequencies of memory CD8 T cell subsets and NKT cells as a percentage of total CD8+ T cells in each cohort. B) CytoGLMM analysis of total CD8 T cell features that are significant predictors of the ELT cohort (left) and the DST cohort (right). Each row represents one marker. The X axis represents log-odds. Bars represent 95% confidence intervals. Markers whose 95% confidence intervals do not cross the red line are considered to be significant predictors of one cohort over the other. C-E) Boxplots showing the median signal intensity (MSI) in naive CD8 T cells (C) or effector memory CD8 T cells (D-E) four markers that were differentially expressed at the patient level between ELT and DST donors. Markers shown in D are expressed at higher levels in ELT patients; markers shown in E are expressed at higher levels in DST patients. Significance values were determined using a Wilcoxon rank-sum test. ns, p > 0.05. *, p < 0.05. **, p < 0.005. ***, p < 0.0005. ****, p < 0.00005.

Whereas naive CD4 T cells differed minimally between DST and ELT donors, naive CD8 T cells had marked differences between the cohorts (**Fig. 6C**). Many more stress-induced molecules, including B7-H6, LFA-3, LLT-1, MICA/B, and Nectin-2 were upregulated in the DST naive CD8 T cells compared to the ELT cohort. Only one marker, CD7, was upregulated on ELT naive CD8 T cells; CD7 expression on CD8 T cells is known to be negatively correlated with HIV response, with high CD7 expression returning upon ART initiation(30) (**Fig. 6C**). By contrast, effector memory CD8 T cells were less altered than effector memory CD4 T cells. CCR2 and CD7 were upregulated in the ELT cohort, along with CD27 and CX3CR1 (**Fig. 6D**). Finally, as observed with nearly every other peripheral immune cell subset, DST cohort CD8 effector memory T cells upregulated the stress - induced proteins LLT-1 and MICA/B (**Fig. 6E**).

## DISCUSSION

In this study, we examined how timing and duration of ART impacts immune composition and phenotype in children living with HIV. We utilized two cohorts of ART-treated CLH from Nairobi, Kenya to compare the immune phenotypes of age-matched CLH who were treated early in life and for a longer duration (median duration 62.3 months) versus those whose treatment was delayed and therefore had shorter-term ART (median duration 8.5 months) at the time of sample collection. Although all CLH were virally suppressed at the time of analysis, we identified striking differences in the peripheral immune systems of early long-term-treated (ELT) children compared to their delayed short-term-treated (DST) counterparts. Overall, we found that the DST children had increased expression of markers of stress and inflammation compared to ELT children. DST children had lower CD4 T cell frequencies and increased abundances of effector memory CD4 and CD8 T cells. Conversely, the ELT children exhibited signs of productive antiviral response, lymphocyte maturity, and resolution of inflammation, along with increased frequencies of CD4 T cells and naive CD4 and CD8 T cells relative to the delayed short-term-treated donors. The high degree of distinction between these two groups of children clearly illustrates the impact of early and longer ART initiation on the pediatric immune system.

The most consistent finding across cell subsets in our analysis is the upregulation of stress-induced markers in the DST cohort. B7-H6, LFA-3, LLT-1, MICA/B, Nectin-2, poliovirus receptor (CD155), and the ULBP proteins, all of which are stress-induced markers recognized by lymphocyte activating receptors, are consistently expressed at higher levels in the monocytes and T cells of delayed short-term-treated donors. The ubiquitous upregulation of these proteins across all peripheral immune cells analyzed suggest that DST children aberrant inflammation compared to ELT children, which is consistent with findings from other studies that have shown that early ART preserves normal immune function in pediatric cohorts(3,31).

NK cells are a key component of the anti-HIV immune response(32,33) and our data demonstrate that there are significant differences between the NK cells of the DST and the ELT cohort. Notably, ELT donor NK cells had an increased frequency of CD56^dim^ CD16^hi^ NK cells, which are classically mature NK cells with a high capacity for direct cytotoxicity and ADCC(34,35). The markers which are significantly associated with the ELT cohort, including CD57, Perforin, CD62L, CD16, and NKp30 also support that these donors have NK cells that are more mature and cytotoxic than those of DST donors. These findings were also mirrored in our unsupervised clustering analysis. The increased abundance of NK cells that appear mature and functional in the ELT cohort compared to the DST cohort suggests that ELT donor NK cells have a higher capacity to mount antiviral responses. This is consistent with the finding that children with delayed ART initiation have impaired NK cell responses(3,31), as do individuals with chronic, untreated HIV(36). Our study is not able to determine whether these alterations in NK cell phenotype are due to the late initiation of ART in the DST cohort or their shorter duration of treatment, but future studies should seek to disentangle these factors to further dissect the dynamics of NK cell suppression and recovery following ART initiation in children.

We observe striking differences in T cell subset frequency between our ELT and DST donors. CD4 T cell frequency, which is lowered in untreated HIV, was significantly lower in the DST donors compared to the ELT donors. We previously found in these cohorts that pediatric patients with delayed onset of ART were unable to fully recover their CD4 T cell counts(37). We also found significant shifts in T cell memory subset distribution in the DST than the ELT cohort: in both the CD4 and CD8 T cell compartments, DST donors had a significant decrease in naive T cell frequency and a corresponding increase in effector memory and effector memory re-expressing CD45RA (EMRA) frequency. This shift was particularly pronounced in CD8 T cells. Loss of naive T cells is a known consequence of untreated HIV-1 infection(38,39) and leaves the host more vulnerable to infection and over-exuberant proinflammatory responses(40). This shift from primarily naive to primarily effector memory T cells in the DST cohort may be caused by the delayed onset of ART, by the persistence of inflammation following untreated HIV infection, shorter duration of ART, or by a combination of these factors.

There were also profound differences in the phenotypes of total, naive, and effector memory CD4 and CD8 T cells between the DST and ELT cohorts. Higher PD-1 expression on the naive and effector memory CD4 T cells of DST donors may impede the accumulation of productive, antigen-specific T cells in response to infection(41). Meanwhile, ELT donor effector memory CD4 T cells exhibited a phenotype that suggests a productive and appropriate antiviral response, with the upregulation of the Th1 markers CCR2 and CCR5(27), CD7(42), and CD127(29). In the CD8 compartment, we found that effector memory CD8 T cells from ELT donors expressed markers that are suggestive of a protective immune response. These include upregulation of CD27, which promotes survival of activated effector memory CD8 T cells(43); CCR2, which regulates trafficking to the site of viral infection(44); CX3CR1, whose increased expression marks more differentiated CD8 T cells with a higher cytotoxic capacity(45); and CD7, whose expression can be downregulated by untreated HIV-1 infection(30). Finally, the effector memory and naive populations along with the total T cell populations in both the CD4 and CD8 compartments of DST donors were marked by the upregulation of stress-induced proteins. Collectively, the differences in composition and phenotypes of both CD4 and CD8 T cells between the DST and ELT cohorts suggest that ELT donors have a T cell pool which is primed for a productive antiviral response, while DST donors have T cells that are stressed, exhausted, and poorly equipped to mount a successful immune response to new infections.

This study provides valuable insight into the effects of ART initiation timing and the dynamics of immune reconstitution in children living with HIV, but nevertheless has limitations that must be considered. We compared age-matched children with differences in time to ART initiation and ART duration because child age profoundly influences immune phenotype. However, this meant that the cohorts used in this study have two major distinguishing factors between them that cannot be disentangled; our cohorts differ in both age at ART initiation and in ART duration. Both of these features influence immune restoration.

The results presented in this study underscore the clinical importance of recognizing the diversity in immune capacity amongst young CLH, which is influenced by duration of ART and age at ART initiation. We examined two pediatric cohorts of the same age with well-suppressed levels of HIV RNA and identified dramatic differences in immune composition and phenotype between the cohorts, with the children who initiated ART later in life and for a shorter duration showing signs of persistent inflammation and immune dysregulation compared to those who were treated soon after birth and for a longer duration. Collectively, these results reinforce that viral suppression by ART in children living with HIV does not return the immune system to a fully “normal” state and earlier longer treatment is required to better reconstitute immune function in CLH.

## MATERIALS & METHODS

### Study Cohorts and Sample Collection

Samples and data used in this study were from two cohorts of pediatric HIV infection conducted in Nairobi, Kenya. Both studies were approved by the University of Washington Institutional Review Board and the Kenyatta National Hospital Ethics and Research Committee. Cohort recruitment and follow-up procedures are described in detail elsewhere(46,47). Briefly, the cohort here referred to as the Early Long-term Treated (ELT) cohort was originally called Optimizing Pediatric HIV Therapy (OPH), NCT00428116, in which HIV-infected, ART-naive infants 1-12 months old were identified at routine HIV-1 testing for prevention of mother-to-child transmission of HIV (PMTCT) clinics and pediatric wards between 2007– 2010. All infants initiated ART after enrollment and were followed for 24 months before being randomized (if eligible) to ART interruption or continued ART(48). In the cohort we refer to here as the Delayed Short-term Treated (DST) cohort [originally called the Pediatric Adherence (PAD) study, NCT00194545], ART-naive children aged 15 months to 12 years who were ART-eligible based on standard of care criteria at the time (WHO disease stage III-IV and/or CD4<15%) were enrolled from Kenyatta National Hospital HIV clinic and pediatric wards between 2004–2007. Children were started on ART and randomized to adherence counseling alone or with a medication diary, then followed monthly for growth, clinical indicators and self-reported adherence.

For both cohorts, blood was collected at 3-6 month intervals during follow-up and was separated into plasma and peripheral blood mononuclear cells (PBMCs). A subset of cohort participants consented and enrolled in extended follow-up with blood collection through up to 8 years on ART. For the sub-study presented here, we restricted analysis to participants who had PBMC samples available that were collected between 4 -8 years of age (to reduce the influence of age-dependent changes in immune profiles) as well as restricted to timepoints when HIV viral load was suppressed below 150 copies/mL during ART at time of CyTOF measurements as well as >3 months prior.

### HIV RNA and DNA quantification

HIV RNA was previously quantified in longitudinal plasma samples from both cohorts using the Gen-Probe HIV RNA assay, lower limit of detection of 150 copies/ml(46,47). Total and intact HIV proviruses were quantified using the cross-subtype intact proviral DNA assay (CS-IPDA) on DNA from cryopreserved PBMCs(49,50). CS-IPDA was performed in triplicate, with additional replicates performed if no intact HIV proviruses were detected in the first 3 replicates, until either intact proviruses were detected or a minimum of 1e5 cells were interrogated. Both total and intact HIV DNA levels are determined for samples with DNA shearing rates of <40% as measured by the RPP30 reference assay(50,51). Data from samples with >40% DNA shearing (n=1) is limited to total HIV DNA because of the impact of shearing on intact HIV DNA quantification. In this analysis, all samples have detectable total HIV DNA. The CS-IPDA is able to detect a single copy of intact HIV DNA(50), and thus, samples with undetectable intact HIV DNA (n=21) were set to 0.5 copies over the number of cells interrogated normalized to 1e6 cells.

### PBMC thawing

PBMC were thawed in a 37C water bath and transferred to RPMI 1640 media supplemented with 10% FBS, 1% L-glutamine, and 1% Penicillin-Streptomycin-Amphotericin B solution (complete medium hereafter referred to as “RP10”). PBMC were counted to determine cell count and viability; any samples with a viability of <50% upon thaw were discarded and not used for CyTOF. 0.5e6 PBMC per sample were set aside in a 37C incubator for later staining, while the rest of the cells were used for NK cell isolation.

### NK cell isolation

NK cell isolation was performed on the remaining PBMC for each donor using the Miltenyi MACS Human NK Cell Isolation Kit. This kit isolates NK cells through magnetic bead-based negative selection. After NK cell isolation, the NK cells were counted and plated in a round-bottom 96-well plate for CyTOF staining. The PBMC that had earlier been set aside in the 37C incubator were also transferred to a separate round-bottom 96-well plate.

### CyTOF staining

The plate containing the isolated NK cells (“NK plate”) and the plate containing the whole PBMC (“PBMC plate”) were centrifuged and all samples were washed once in 1X PBS. The samples were then stained with a Cisplatin-based viability stain at a Cisplatin concentration of 25 uM for 60 seconds; the stain was subsequently quenched with the 1:1 addition of undiluted fetal bovine serum. The samples were washed twice with CyFACS buffer (1X PBS, 0.1% BSA, 2 mM EDTA, 0.05% sodium azide) and then stained for 30 minutes with a Palladium-CD45 barcoding scheme as previously described(16). After barcode staining, the samples were washed three times with CyFACS buffer to ensure complete removal of any unbound CD45 antibody and combined into sets of barcodes, hereafter referred to as “barcoded samples”. These barcoded samples were then stained with the surface NK cell panel (NK cell plate) or the surface PBMC panel (PBMC plate) (Table X). All panels were prepared in advance and lyophilized or frozen in aliquots at -80C to ensure consistency between batches. The stained barcoded samples were washed again and fixed in 2% paraformaldehyde (PFA), then permeabilized. The NK cell plate was then stained with the intracellular NK panel (Table X). Finally, the samples were washed thrice more in CyFACS buffer and resuspended in CyPBS supplemented with 2% PFA and an iridium DNA intercalator and stored at 4C until collection. Data were collected on a Helios mass cytometer. Before collection, samples were washed with CyFACS buffer and Milli-Q water before being resuspended in 1× EQ beads (Fluidigm) for collection.

### CyTOF data pre-processing

Prior to analysis, FCS files were normalized and debarcoded using the functions of the same names in the *Premessa* package(52,53). The normalized and debarcoded FCS files were then transferred to FlowJo v10.9.0, which was used to manually remove EQ beads, doublets, dead cells, and debris from all samples. Any contaminating non-NK cells were manually gated out of the NK cell samples (**Fig. S1-2**). The whole PBMC files were manually gated into major immune cell subsets based on expression of lineage markers; T cells were further divided into memory subsets (**Fig. S1**). The FCS files for each cell subset (monocytes, B cells, total CD4 T cells, total CD8 T cells, NK cells, naive CD4 T cells, naive CD8 T cells, effector memory CD4 T cells, and effector memory CD8 T cells) were then exported from FlowJo for further analysis.

### CyTOF data analysis

The FCS files exported from FlowJo were imported into Rstudio using the *FlowCore* package. Mean signal intensity (MSI) values were arcsinh transformed (cofactor = 5) in order to account for heteroskedasticity in the data. **<u>MU</u>**ltivariate modeling with minimally biased **<u>V</u>**ariable selection in **<u>R</u>** (MUVR) analysis was performed on the samples using the *MUVR* package(17) to identify the key variables in distinguishing between our two cohorts and a generalized linear model (GLM) with the Benjamini-Hochberg correction for multiple hypothesis testing was then used to identify the correlation between each of the variables selected by MUVR and the binary outcome variable (ELT vs. DST).

Unsupervised analyses of NK cells and monocytes in our dataset were performed by first coercing the relevant FCS files into Seurat objects using the *Seurat* package(54,55). Unsupervised clustering was performed using the Phenotyping by Accelerated Refined Community-partitioning (PARC) algorithm(21). Clustering was performed at multiple different resolutions and we selected a resolution to move forward with by plotting cluster sizes and relationships using the *clustree* package and choosing the resolution at which the cluster identities became relatively stable. UMAP embeddings of the arcsinh transformed data were generated using the *uwot* package; all markers were used in the generation of the UMAP.

The *CytoGLMM* package was used to identify markers within each cell type that were significantly associated with one cohort over the other. This method is described in more detail elsewhere(19).

### Data visualization

Boxplots showing the cell subset frequencies and untransformed MSI values of various markers were generated using the *ggplot2* package. Colors were generated using the *MetBrewer* package or selected manually. UMAPs were plotted using *Seurat*. Heatmaps were generated using the *ComplexHeatmap* package.

## DATA AND CODE AVAILABILITY

The normalized and debarcoded .fcs files for all data used in this study along with their accompanying metadata are available on Immport under accession number SDY2750. The code used for analysis in this study is available at https://github.com/BlishLab/pediatric_hiv_cytof.

## ACKNOWLEDGEMENTS

We thank the children and parents who participated in this study, and the team of researchers at University of Nairobi and Kenyatta National Hospital, including clinicians, laboratory technicians and managers, data managers and peer-mentors. We would like to thank the Stanford Human Immune Monitoring Core (HIMC) for their assistance with the collection of CyTOF data. Thank you to Dr. Geoff Ivison, whose code for formatting CyTOF data for statistical analysis in R was adapted for use in this project. We thank Dr. Aaron Wilk for creating the pipeline used in this manuscript for the unsupervised analysis of CyTOF data. Finally, we thank Dr. Matthew Kaufmann and Minne Lee for their insightful commentary on this work.

## SUPPLEMENTAL MATERIAL CAPTIONS

**Supplementary Figure 1: Gating strategy for identifying major immune cell subsets within whole PBMC CyTOF data.** Example flow plots showing the gating schema used to identify major cell subsets in whole PBMC CyTOF data. A) Gating on normalized, debarcoded .fcs files to remove QC beads, doublets, debris, and dead cells. B) Gating on the live, intact cell population derived in (A) to identify major immune cell subsets for the plots shown in Fig. 2. Arrows indicate downstream gates.

**Supplementary Figure 2: Gating strategy for eliminating contaminating non-NK cells within purified NK cells CyTOF data.** Example flow plots showing the gating schema used to identify NK cells and NK cell subsets in purified NK cell CyTOF data. A) Gating on normalized, debarcoded .fcs files to remove QC beads, doublets, debris, and dead cells. B) Gating on the live, intact cell population derived in (A) to remove any contaminating non-NK cells. LILRB1/CD56 and HLA-DR/CD56 gates were used to remove CD16+ CD14-monocytes, which express high levels of these markers and are frequently accidentally included in CD56-CD16+ NK cell gates. Arrows indicate downstream gates.

**Supplementary Figure 3: Gating strategy for identifying memory T cell subsets within whole PBMC CyTOF data.** Example flow plots showing the gating schema used to identify memory T cell subsets in CyTOF data. The first plot (center) shows live, intact, CD3+, CD8+, CD4-, CD56-cells derived from whole PBMC. Arrows indicate downstream gates.

**Supplementary Table 1: Markers used in MUVR analysis.**

